# Bootstrap-based criteria for identifying differences between learned Bayesian networks

**DOI:** 10.1101/2025.11.04.686685

**Authors:** Rosie Berners-Lee, V Anne Smith

**Author notes:** Address: School of Biology, Sir Harold Mitchell Building, Greenside Place, St Andrews, KY16 9TH, UK.

## Abstract

Bayesian networks provide a powerful framework for learning dependencies from data, and they are widely used to probe structure in biological systems. Biological systems are governed by complex networks of interactions, and uncovering these interactions and comparing them across conditions is central to understanding biological mechanisms. However, when comparing Bayesian networks, it can be difficult to determine whether observed differences are substantial enough to reflect genuine differences in the underlying systems generating the data. Here, we address this by developing bootstrap-based criteria for identifying such differences and demonstrate their performance using simulated data from synthetic Bayesian networks. Both edge-level and whole-network connectivity comparisons reliably identified when underlying networks differed, even when this involved only 5% of edges, while distinguishing these differences from sampling variation. However, even with large datasets, the criteria were unable to recover specific edge differences. Thus distinguishing that networks differed was possible, but not the specific ways they differed. These criteria establish a framework for more robust and standardised Bayesian network comparisons, with broad potential for real-world applications.

## Introduction

### Introduction to Bayesian networks

The complexity of biological systems arises from the intricate networks of interactions that govern them. Understanding these networks, and how they change across conditions, is crucial for uncovering the mechanisms that drive biological processes. Bayesian networks (BNs) provide a powerful framework for modelling these interactions as statistical dependencies [1]. The statistical dependencies represented in a BN can be learned from observational data [2]. However, when comparing BNs, it can be difficult to determine whether observed differences are substantial enough to reflect true differences between the underlying systems [3, 4]. This highlights the need for robust criteria to assess the significance of structural differences between BNs, which this study aims to address. To approach this, it is necessary to first outline what BNs are and how they are learned, before reviewing existing approaches for comparing network structures more broadly.

A BN is a directed acyclic graph (DAG), where each node represents a variable and directed edges indicate direct dependencies between them. Since BNs are acyclic, they contain no loops, meaning that no variable can directly or indirectly influence itself. Relationships between nodes are described using familial terminology, with edges directed from “parent” to “child” nodes. The edges encode a minimal set of conditional independencies, such that each node is conditionally independent of its non-descendants, given its parents (Fig 1A) [5]. Each node in a BN is associated with parameters that define the probabilities of the node being in each state, given every possible combination of its parents’ states. Therefore, a BN consists of two components: its structure and its sets of parameters corresponding to each node. A widely used type of BNs are discrete BNs, where each variable can only take on a finite set of discrete states and each node’s parameters are represented by a Conditional Probability Table (CPT), where each row corresponds to a unique parent state combination [5].

**Fig 1.**
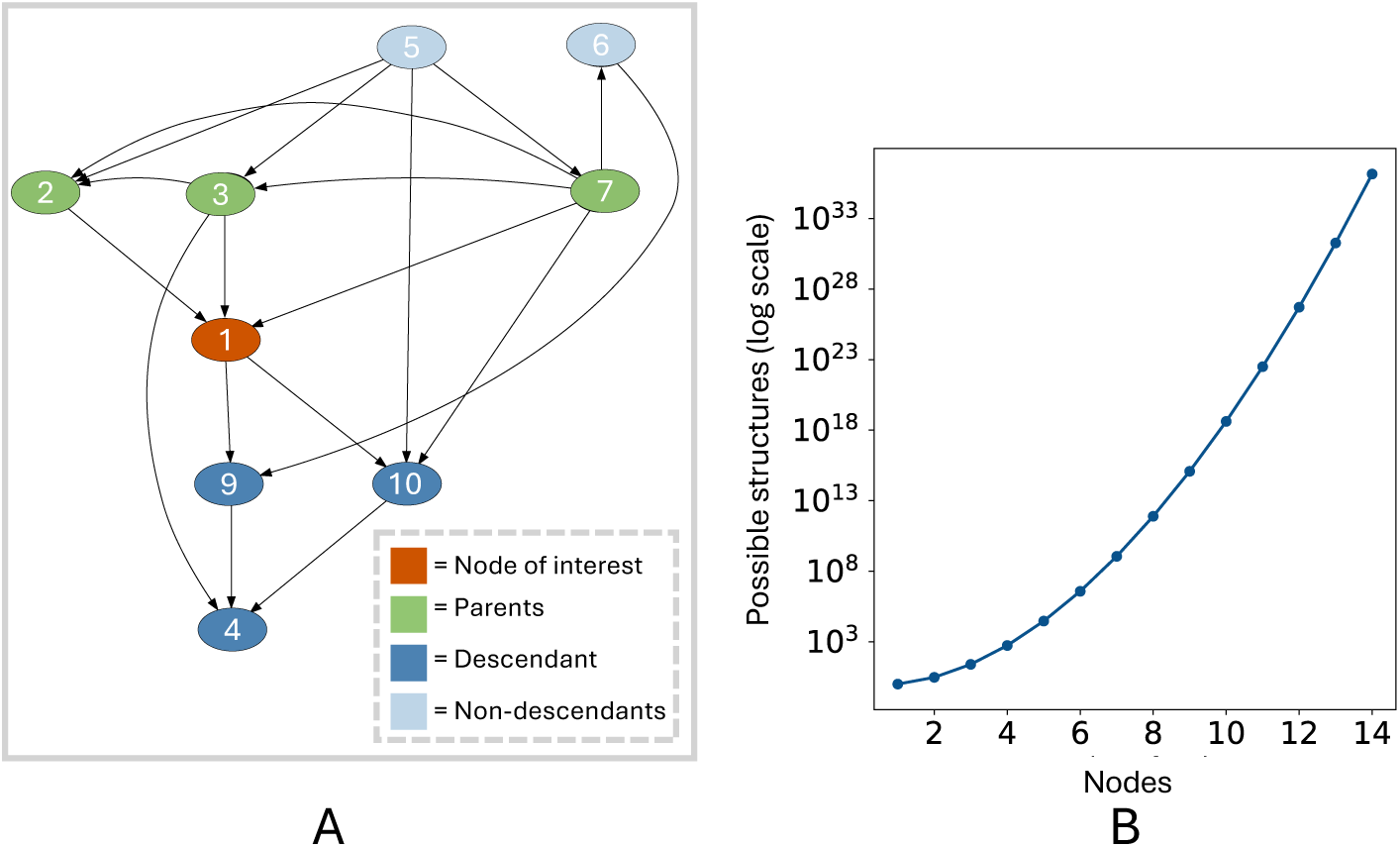
Bayesian network Example. A: Node-edge diagram representation of an example 10-node Bayesian network (BN). Nodes (*ovals*) represent variables and edges (*arrows*) represent direct dependencies between them such that Node 1 (*orange*) is conditionally independent of all its non-descendants (*light blue*) given its parents (*green*). B: Super-exponential increase in the number of possible BN structures with increasing numbers of nodes.

### Heuristic searches

A popular method of learning BNs from data is via search-and-score, where a search algorithm optimises a score to find BNs that fit the data. Bayesian Scoring Metrics (BSMs) are used to assess how effectively a BN explains the data it represents whilst penalising for complexity to avoid overfitting. However, the number of possible BN structures increases super-exponentially with nodes (Fig 1B), making it computationally infeasible to exhaustively evaluate all structures to determine which has the best score [3, 6]. Heuristic search algorithms address this by exploring subsets of networks to identify high-scoring structures that fit the data well enough. These algorithms work by iterating through cycles of scoring a network, modifying it, re-scoring the new structure, and deciding whether to accept the change [1, 5]. For more information, see S1 Appendix.

The stochastic nature of heuristic searches means different search runs often yield varying results, as multiple structures can achieve similar high scores [3, 4]. Therefore, to ensure reproducibility, a search procedure should be established that produces consistent BN structures. Since achieving exact reproducibility is often infeasible, instead, the procedure should aim for sufficient consistency for the resulting BNs to support the same conclusions [4]. Search procedures often combine multiple high-scoring BNs from the same or different search runs to build a consensus network [7, 8]. They can also be used to estimate the probabilities of specific edges or other network features [3, 9].

### Comparing networks

BNs are increasingly being used in diverse fields of biology, including molecular biology, neuroscience, and ecology, to infer underlying regulatory interactions from noisy high-dimensional data [10]. Their ability to capture conditional dependencies and handle sparse datasets makes them especially suitable for reconstructing gene regulatory networks (GRNs), including in non-model organisms where prior knowledge is limited. In biological research, comparing BNs across conditions has helped identify novel drug targets by revealing changes in GRNs in response to disease or treatment [11–14]. More generally, comparing BNs is valuable for understanding how different systems function or how the same system responds to perturbations. However, since multiple BN structures can explain the same data equally well, non-identical BNs do not necessarily reflect genuine differences in the underlying processes [3, 4]. Currently, these differences are often assessed on an ad hoc basis, which is time-consuming, hinders cross-study comparisons, and undermines confidence in conclusions. This highlights the need for robust, standardised criteria to determine when differences between BNs are substantial enough to infer that the underlying processes they represent genuinely differ. Despite their growing use, this aspect of BN research remains largely unexplored.

Existing methods for comparing network structures, although not specific to BNs, offer valuable approaches for addressing this gap [15]. A commonly used measure in graph theory is graph edit distance, which counts the number of alterations required to transform one graph into another [16]. While effective in fields like graphics and pattern analysis, it is less suitable for comparing complex natural systems [17]. For networks with known node correspondence, as is common with BNs, the simplest approach is to compare edge presence directly. While rarely used alone, this forms the basis of more advanced methods, such as the DeltaCon similarity score [15]. DeltaCon compares global connectivity by combining information on edge presence with the number of edges connected to each node (its degree). It incorporates all possible paths (sequences of edges) between variables, weighting shorter paths more heavily to reflect stronger, more direct connections [18, 19]. This has been applied to diverse topics, including brain connectivity and email communication networks [19].

An alternative measure is cut distance, which partitions each network into the same two sets of nodes and measures the total number (or weight) of edges going from one set to the other. The cut distance is the maximum difference in this between the two networks across all possible partitions [20]. Both cut distance and DeltaCon can be applied to directed and weighted networks, making them suitable for diverse network types, including BNs. However, by focusing only on the partition with the largest difference, cut distance is highly sensitive to local changes. Its reliance on evaluating many possible partitions also makes it computationally intensive, often making DeltaCon the more practical choice [15].

For more specific comparisons of node-level structure, networks can also be compared using centrality measures [17]. Centrality describes a measurable feature of a node reflecting its role or position in the network. For example, degree centrality captures the number of connections per node, closeness centrality measures proximity to other nodes via shortest paths, and betweenness centrality reflects how often a node lies on the shortest paths between other nodes. Centrality distance sums the differences in centrality values across corresponding nodes in the two networks, and the choice of centrality depends on the focus of the analysis [17], making it less relevant for a generic methodology.

In this paper, we focus on using direct comparisons of edge presence for a fine-grained measure and the DeltaCon similarity score for a more granular measure of whole-network connectivity.

### Aims and methodological approach

Here, we develop statistical criteria for determining whether differences between BNs are substantial enough to conclude that the underlying systems they represent differ. To do this, datasets were simulated from pairs of randomly generated synthetic BNs, to represent the two datasets in a comparison. We used a bootstrap-based method to generate distributions associated with each dataset, which can then be statistically compared. Each dataset was bootstrapped multiple times, and BN structures were inferred from each bootstrapped dataset, producing sets of bootstrap-derived BNs (BDBNs). Differences were then assessed by comparing edge frequencies and whole-network connectivity across BDBNs corresponding to either the same or different synthetic BNs.

Using simulated data from synthetic BNs provides a valuable testbed for method development since, unlike in real-world scenarios, the ground truth is known, enabling the criteria to be validated [2]. Bootstrapping amplifies the dataset and aims to mimic variability across independent replicates, helping ensure the criteria remain robust when data is limited [21]. We show that, although unable to recover specific edge differences, comparisons of both edge frequencies and whole-network connectivity are able to distinguish true structural differences between underlying systems from spurious differences introduced by dataset variation. These approaches provide a valuable frameworks for comparing BNs across diverse fields.

## Materials and methods

### Software and processes

BN structure learning was performed using Banjo (version 2.2) [8]. Custom R scripts (version 4.4.2) were used to generate synthetic BNs and datasets. All other analyses, including workflow automation, data preprocessing, and evaluation of model outputs, were conducted using custom Python scripts (version 3.6.8). Code to reproduce results and apply the criteria to data is available at https://github.com/rb2065/comparing_BNs.git.

### Synthetic networks and datasets

#### Generating synthetic Bayesian networks

Random BNs were generated using the random.graph function from R’s bnlearn package. A network size of 40 nodes was chosen as it is near the practical upper limit where heuristic searches can still explore a sufficiently large subset of the search space to identify high-scoring structures within a reasonable time. The maximum number of parents per node was three. Ten random BNs were generated, and from each, three additional BNs were generated by removing 50%, 25% or 5% of the edges and then attempting to reintroduce an equivalent proportion. Reintroduced edges could not include previously removed edges or exceed the three-parent limit, and cycle checking was applied to ensure the networks remained DAGs. Due to these constraints, the final networks did not necessarily retain the original edge count. This resulted in pairs of synthetic BNs differing in approximately 50%, 25%, and 5% of their edges, referred to as the BN50, BN25, and BN5 pairs, respectively. These varying levels of structural difference were used to assess the sensitivity of the comparison criteria to progressively smaller differences between underlying networks. Unless otherwise stated, and excluding analyses for search procedure optimisation (see S3 Appendix), results are averaged over the 10 independent repeats.

#### Simulating data and bootstrapping

For each BN, random CPTs for each node were generated using the rdirichlet function from R’s MCMpack package. Each row in a CPT corresponds to a unique parent state combination, columns to the node’s possible states, and values to their conditional probabilities given the parent state combination [5]. For simplicity, each node was restricted to two possible states. The CPTs were calculated using a Dirichlet distribution to ensure that the total probabilities in each row sum to one. The concentration parameter (*α*) of the Dirichlet distribution controls the spread of the probability distribution [22]. To ensure that subsequently simulated data reflected the dependencies encoded in the BN structures, *α* values were chosen based on node parentage: *α* = 5 for nodes without parents produced more uniform probabilities across states, avoiding bias, while *α* = 0.5 for nodes with parents encouraged more extreme probabilities, thereby strengthening the influence of parent nodes on their children’s states. Two independent synthetic datasets of 40 samples were simulated from each BN structure and corresponding CPTs using bnlearn’s rbn function. This small dataset size was chosen to reflect the limited sample numbers typically available in GRN studies, particularly in non-model organisms. These studies are a valuable and increasingly common application of BNs in biology. Therefore, using a similarly constrained dataset tests the criteria under realistic conditions while also providing a stringent case that increases confidence in their performance on larger datasets. Additional larger datasets of 400 and 4000 samples were generated for one BN50 and one BN25 example for use in a single analysis.

#### Bootstrapping

Each synthetic dataset was bootstrapped (sampled with replacement) to generate 100 bootstrap-derived datasets of the same size as the original. Using the search procedure described subsequently, for each bootstrap-derived dataset, the probability of each possible undirected edge was determined and an edge probability threshold was applied, generating two sets of 100 BDBNs corresponding to each synthetic BN.

### Establishing a heuristic search procedure

#### Software and settings for heuristic searches

Heuristic searches were performed on each bootstrapped dataset using Banjo software. Banjo uses the Bayesian Dirichlet equivalent (BDe) BSM to identify top networks and can compute a consensus network from the top *n* networks, weighted by score [8]. To reduce the search space, for all searches, the parent count per node was capped at three [23]. To allow broader exploration of the search space, all searches used a network modification stage that applied a single random local move per step, rather than selecting the highest-scoring option from all possible local moves. Based on search optimisation (see S3 Appendix), to balance search consistency and network recoverability with run-time, a stopping criterion of 10 random restarts was applied, such that each top network was the highest-scoring among 10 local maxima.

#### Network averaging search procedure

Since initial explorations revealed highly complex search spaces (see S2 Appendix), a network averaging approach, applied over 100 greedy searches (each with 10 random restarts), was used, inspired by [7] (S2 Fig). For this, the top network from each search was converted to an adjacency matrix (**A**) where *A_i,j_* = 1 if an edge existed between nodes *i* and *j*, and *A_i,j_* = 0 otherwise. To increase reproducibility, edge directions were ignored. Since this study focuses on static networks and the primary interest is identifying differences in connectivity patterns rather than inferring directionality of interactions, omitting edge directionality has minimal impact on interpretation.

The resulting 100 adjacency matrices were used to construct a difference matrix (**M**) where *M_i,j_* represents the sum of the absolute differences between the adjacency matrices of networks *i* and *j*. Hierarchical clustering was applied to **M** to identify groups of similar networks. From this, a similarity matrix (**K**) was computed where *K_i,j_* represents the similarity between networks *i* and *j*, normalised to between 0 and 1.

Weights for each network were derived from their relative BDe scores, extracted from Banjo search reports, and used to construct a weight matrix (**W**), where *W_i,j_* is the product of the weights of networks *i* and *j*. A covariance matrix (Σ) was then computed as the element-wise product of the similarity and weights matrices.

The covariance matrix was used to fit an intercept-only generalized least squares (GLS) model for each possible edge, with the dependent variable representing the presence or absence of the edge across networks. This approach weighted each network’s influence based on both its relative score and similarity to other networks, giving greater influence to higher-scoring networks and those more similar to others. The model outputs represented the probability of each edge being present in the underlying network. A minimum edge probability threshold (0.75 unless otherwise stated) was then applied to determine which edges would be included in the final BN (See S3 Appendix).

### Approaches to comparing bootstrap-derived Bayesian networks

To assess whether significant differences were more prevalent when comparing sets of BDBNs from different underlying networks (between-BN; B-BN) versus the same underlying network (within-BN; W-BN), each BDBN set was randomly split in half. Three comparison types were used: B-BN comparisons, where half-sets came from different underlying synthetic networks; same-dataset within-BN (SD-W-BN) comparisons, where both half-sets were from the same BDBN set; and across-dataset within-BN (AD-W-BN) comparisons, where half-sets came from separate datasets independently simulated from the same synthetic BN (Fig 2).

**Fig 2.**
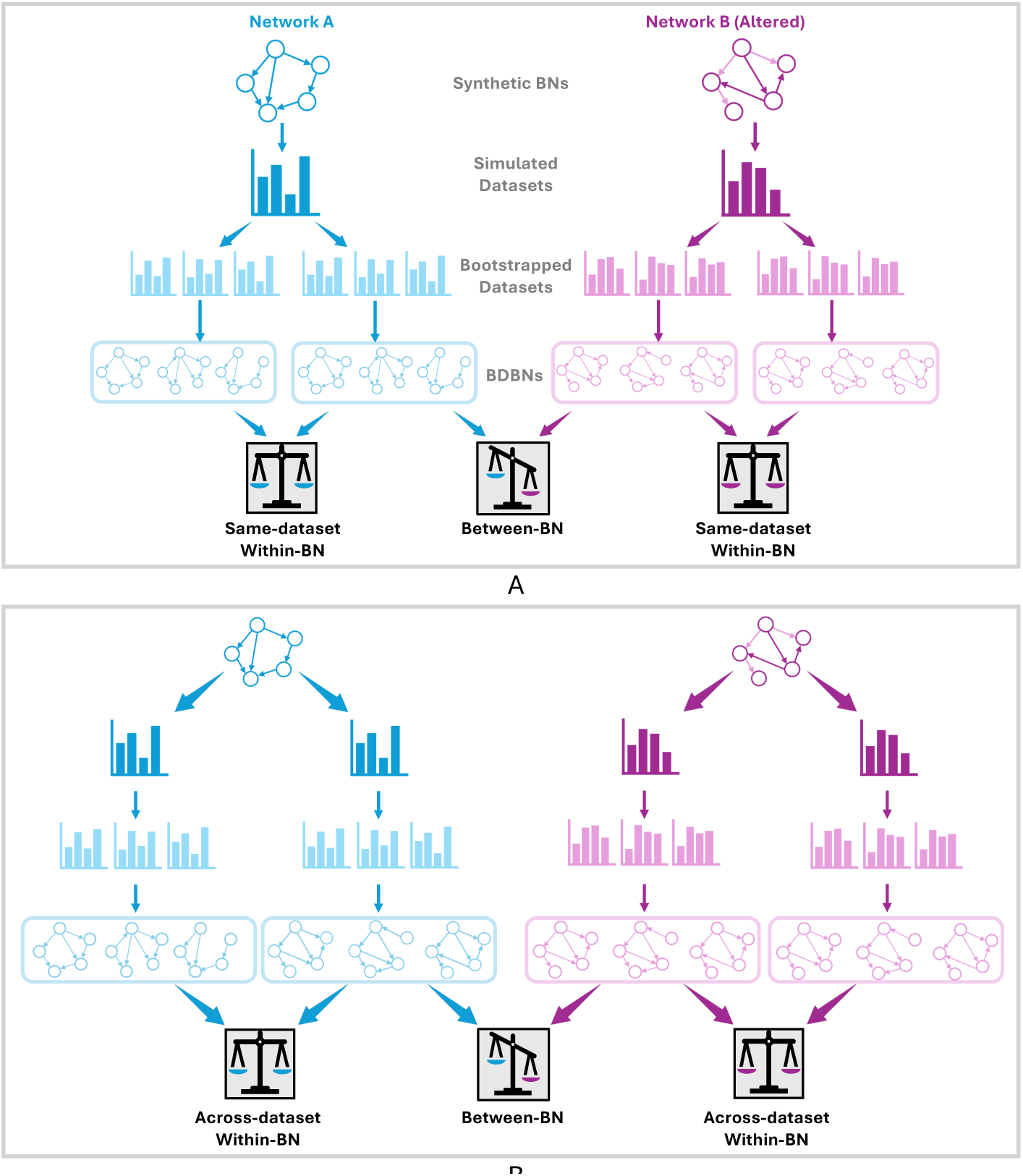
Types of network comparisons. Workflows for comparing sets of bootstrap-derived Bayesian networks (BDBNs) from the same (within-BN) or different (between-BN) underlying networks. Datasets are simulated from pairs of random synthetic Bayesian networks (BNs) differing by a fixed proportion of edges. Bootstrap resampling is applied to produce groups of bootstrapped datasets, and a BDBN is learned from each. Comparisons are conducted between groups of BDBNs corresponding to either the same synthetic BN (within-BN) or different synthetic BNs (between-BN). A: Same-dataset within-BN comparisons: one dataset is simulated from each synthetic BN and bootstrapped to produce two groups of BDBNs per network. B: Across-dataset within-BN comparisons: two independent datasets are simulated from each synthetic BN, and bootstrapping is applied separately to produce two groups of BDBNs per network.

For each comparison type, edge frequencies and DeltaCon scores were compared between the two half-sets. Comparisons were repeated 100 times with different random splits of the BDBN sets, generating a distribution of 100 comparison values for each type. Mean values were compared using the permutation test with 10,000 permutations, with the Bonferroni correction [24] applied to an initial significance threshold of *α* = 0.05 (divided by 2 to account for testing that both SD-W-BN and AD-W-BN comparisons were significantly less different than B-BN comparisons). The difference criteria were considered reliable if significant differences were detected for B-BN but not for either W-BN comparison. Significant differences for AD-W-BN comparisons would indicate that the criterion could not fully distinguish true network differences from variation between independent datasets.

### Comparing edge frequencies across networks

#### Identifying significant differences in edge frequencies

For each comparison type, the frequency of each possible edge was compared between sets of BDBNs and significant differences in edge frequencies were identified using Fisher’s Exact test. Only edges present in at least one set were compared and, to account for multiple testing, the significance threshold of *α* = 0.05 was adjusted using the Bonferroni correction [24], calculated as 0.05 divided by the number of edges compared. The number of edges found to be significantly different was used as the comparison value to assesses network differences. The underlying BNs were considered significantly different if the mean number of significant differences in edge frequencies from B-BN comparisons was significantly higher than both SD-W-BN and AD-W-BN comparisons. Edge-frequency comparisons were repeated for BDBN sets of varying sizes (20–100) to determine the minimum set size required to detect significant differences.

#### Recovery of underlying edge differences

To assess how well the BDBN edge comparison approach detects true differences between the underlying networks, edges with significantly different frequencies between pairs of BDBN sets were compared to the actual edge differences between the corresponding synthetic networks. This analysis was conducted only for the BN25 and BN50 pairs. TPs, FPs and FNs were defined as edges that were correctly identified as different, incorrectly identified as different, or incorrectly identified as not different, respectively. From these definitions, precision and recall (Eq 1,Eq2), along with TP and FP counts, were calculated across different minimum edge probability thresholds (0.25, 0.5 and 0.75) and numbers of bootstraps (20–500) to assess their impact. For a single BN50 and BN25 example, the recovery of underlying differences was also assessed using BDBNs learned from larger datasets of 400 and 4000 samples.

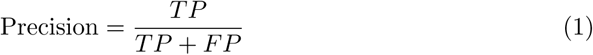

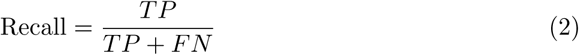

### Comparing whole-network connectivity across networks

To compare pairs of BN structures more globally, whole-network similarity was assessed using the DeltaCon algorithm [18]. DeltaCon similarity scores were calculated between all possible pairs of individual BDBNs from the half-sets being compared, producing a distribution of scores for each comparison type. To calculate the DeltaCon similarity score, for each BDBN in the pair, an adjacency matrix (**A**) was constructed, as previously defined. From this, a degree matrix (**D**) was computed by summing each row of **A**, such that *D_i,i_* represents the number of edges connected to node *i* (its degree), and all off-diagonal elements are zero. An affinity matrix (**S**) was then computed as:

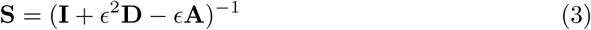

Where **I** denotes the identity matrix, a square matrix with ones on the main diagonal and zeros elsewhere. Each element *S_i,j_* represents the influence of node *i* on node *j*, incorporating both direct and indirect connections. Eq 3 expands as a power series where paths of length *k* contribute with a weight proportional to *ɛ^k^*. As recommended by [18], *ɛ* was set as:

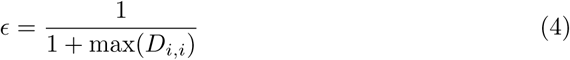

Since *ɛ <* 1, longer paths are weighted less, ensuring that influence between nodes reduces with increasing path length. The DeltaCon similarity score was then calculated by first computing the DeltaCon distance (*δ*) as:

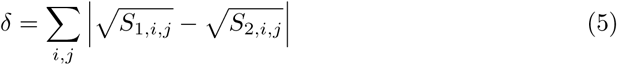

Here, square rooting reduces the impact of large differences, enhancing sensitivity to smaller structural changes. Finally, the DelaCon similarity scores (*sim*) were calculated by normalising *δ* to between 1 and 0 using:

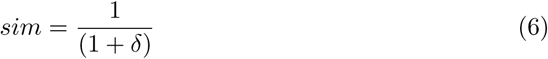

Such that for identical networks, *sim* = 1, and *sim* → 0 as structural differences increase.

The underlying BNs were considered significantly different if the mean B-BN similarity score was significantly lower than both W-BN means.

### Visualising Bayesian network comparisons

While node-edge diagrams (Fig 3A) are the most common way to visualise BNs and are intuitive for representing small structures, they quickly become cluttered and difficult to interpret as network size and complexity increase, especially when comparing multiple networks [25–27]. To support interpretation and comparison of network structures, this study instead used matrix-based visualisation throughout. This approach is more scalable, with rows and columns representing child and parent nodes, respectively, and cell values indicating edge presence (Fig 3B). Matrix-based visualisation enables clear comparison between networks, as cell values can indicate whether edges are shared across conditions or consistently recovered across repeated searches [26]. Such differences can be visualised using variations in colour or intensity. Notably, matrices representing undirected networks are symmetric about the main diagonal (Fig 3C). This allows results from different analyses to be displayed on each triangular half, further facilitating comparison.

**Fig 3.**
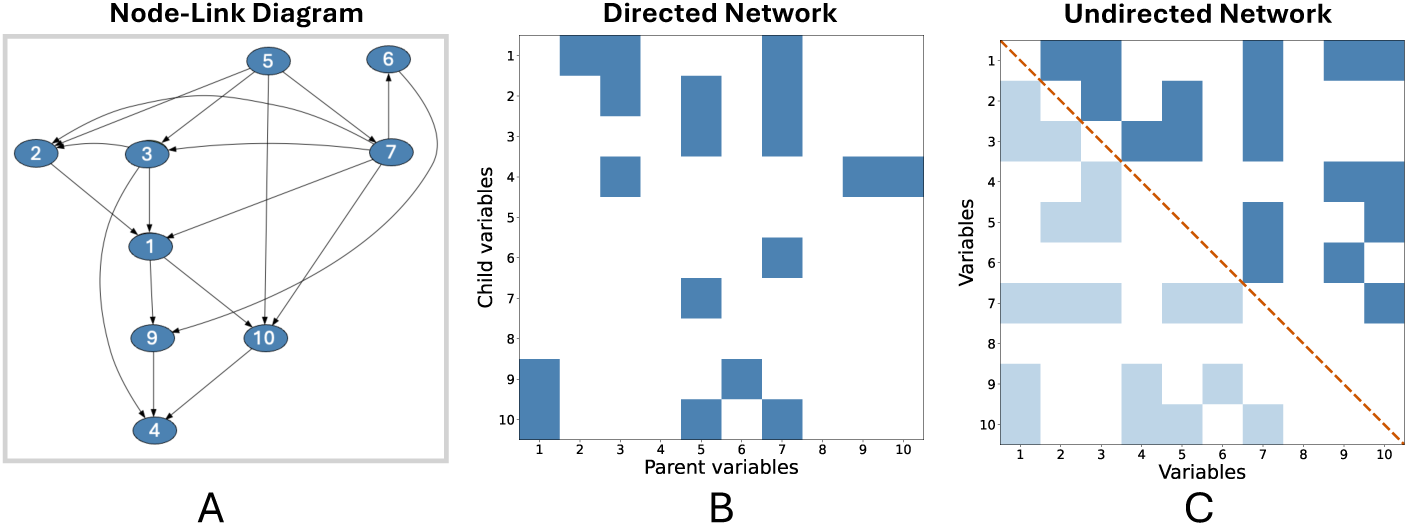
Matrix-based Bayesian network visualisation. A: Node-edge diagram representation of an example 10-node Bayesian network (BN). Nodes (*ovals*) represent variables and edges (*arrows*) represent direct dependencies between them. B: Matrix representation of the BN in A, where *columns* represent parent variables, *rows* represent child variables, and *coloured cells* indicate edges between them. C: Undirected version of the network in B, resulting in symmetry along the main diagonal (*orange line*). This makes half of the matrix redundant, allowing it to be repurposed to display an alternative network or analysis for easier comparison.

## Results

### Recovery of underlying individual edge differences

Sets of 100 BDBNs were generated for the BN50 and BN25 synthetic BN pairs, and edge frequencies were compared between sets to assess whether true underlying edge differences could be recovered. Although consistency within each BDBN set was lower than across repeated searches on the same dataset, edge probabilities remained similar within sets, and visual inspection revealed clear edge differences between the BDBN sets corresponding to each network in both BN pairs (S4 Fig).

Despite these qualitative differences, recovery of true structural differences was poor. Only 2.14±0.90 TPs were identified for the BN50 pair, and 1.71±1.50 for BN25 (Figs 4A-4C), resulting in low precision and recall. Given this poor difference recovery even for networks with larger structural differences, the BN5 pair — which differed by only a small proportion of edges — was excluded from this analysis. Lowering the edge probability threshold increased recall due to the larger number of significant edge differences identified (Fig 4D). However, precision remained low, indicating that adjusting the edge probability threshold would not substantially improve precision. The recovery of true edge differences did not improve when the dataset size was increased to 400 or 4000 samples (S5 Fig, S6 Fig).

**Fig 4.**
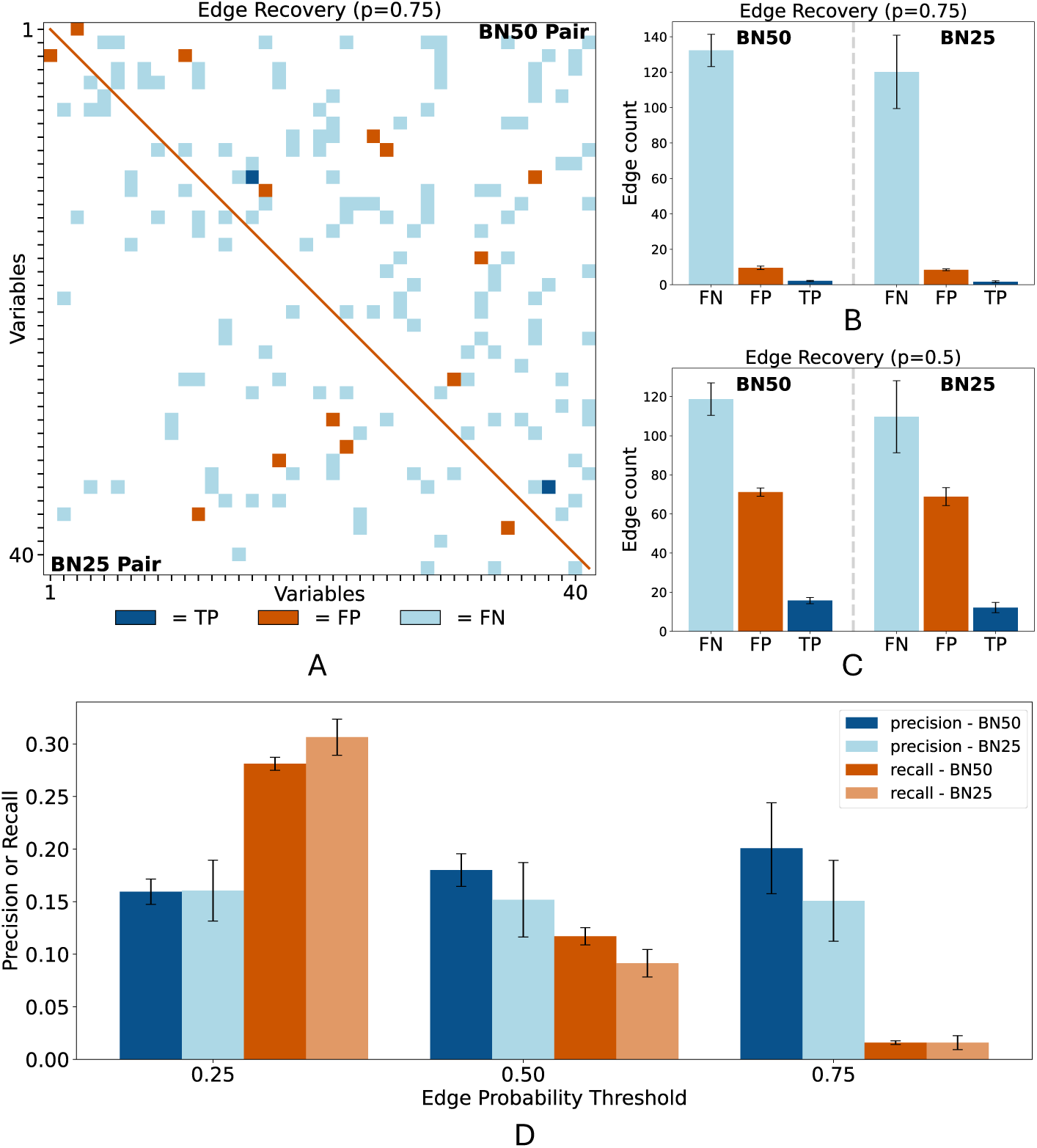
Recovery of underlying differences between synthetic Bayesian networks. A: Example of recovered differences between the BN50 (*top right*) and BN25 (*bottom left*) network pairs after applying a minimum edge probability threshold of 0.75. True positives (TP, *dark blue*), false positives (FP, *orange*), and false negatives (FN, *light blue*) represent edges correctly identified as different, incorrectly identified as different, and incorrectly identified as not different, respectively. B,C: TP (*dark blue*), FP (*orange*), and FN (*light blue*) counts for BN50 (*left*) and BN25 (*right*) after applying minimum edge probability thresholds of 0.75 (B) or 0.5 (C). D: Precision (*blue*) and recall (*orange*) of edge difference recovery across minimum edge probability thresholds. Results for BN50 and BN25 are shown in *darker* and *lighter* shades, respectively. Bars in (B-D) show mean values, with error bars representing ±1 standard error across 10 repeats.

To further investigate the poor recovery of true edge differences, for a single BN50 example, the number of BDBNs used in the comparison was varied from 20 to 500 to assess its effect on precision (S6 FigA). For edge probability thresholds of 0.75 and 0.5, precision initially spiked at low BDBN counts but quickly declined and plateaued. This early spike was driven by the small number of differences detected, where TPs made up a higher proportion. While this suggests some true differences may have stronger effects than spurious ones, the low absolute number of TPs made reduced BDBNs an unviable solution. At a 0.9 threshold, no edge differences were identified until ∼ 100 BDBNs, where a single TP caused 100% precision which decreased when FPs were identified at ∼ 400 BDBNs.

Across all three thresholds, both TPs and FPs increased with increasing BDBNs, though the effect was more pronounced at lower probability thresholds (S7 FigB-C). However, because FPs consistently outnumbered TPs, overall precision remained low. In real-world settings with unknown ground truth, TPs would be indistinguishable from FPs. Therefore, given the goal of identifying a meaningful number of true differences and the computational cost of generating more BDBNs, adjusting the threshold or number of BDBNs alone is unlikely to resolve the poor recovery performance. As a result, the threshold was kept at 0.75 and the number of BDBNs at 100, but the focus shifted away from identifying individual true differences and toward assessing whether the underlying networks differed overall.

### Network differences based on edge frequency comparisons

Although most significant differences in edge frequencies identified did not correspond to the true underlying differences, they may still indirectly reflect structural variation between networks. If so, comparisons between BDBN sets derived from different underlying networks (between-BN; B-BN) should yield significantly more edge differences than those from the same network (within-BN; W-BN). Indeed, all BN50, BN25, and BN5 B-BN comparisons of 100 BDBNs showed significantly more edge differences (*p* = 0.0002) than same-dataset W-BN (SD-W-BN) comparisons, where none were detected (Figs 5A-C).

**Fig 5.**
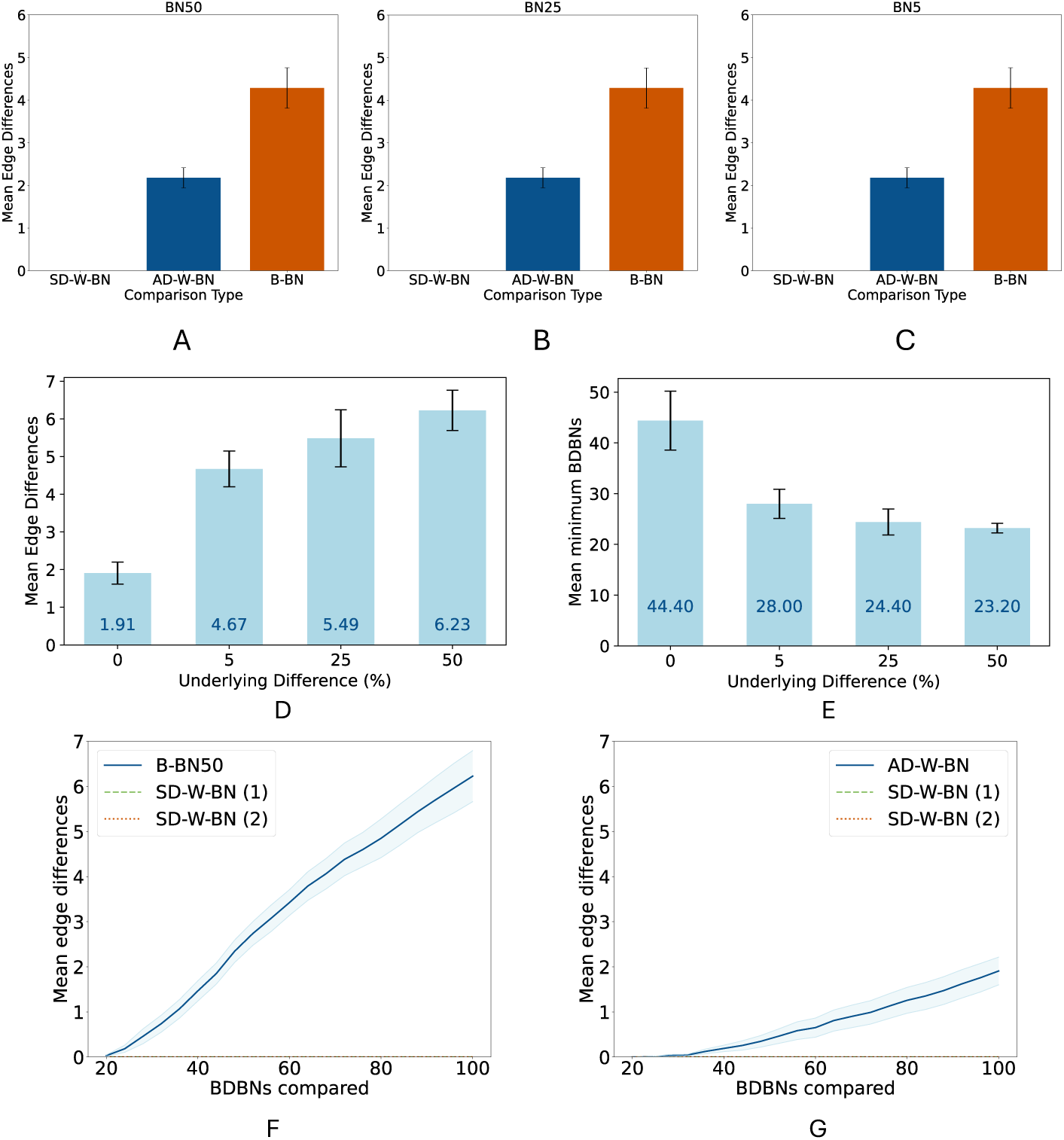
Edge frequency comparisons between sets of bootstrap-derived Bayesian network (BDBN) A-C: Mean number of significant differences in edge frequencies identified between sets of 100 BDBNs from across-dataset within-BN (AD-W-BN, *dark blue*), and between-BN (B-BN, *orange*) comparisons. Same-dataset within-BN comparisons (SD-W-BN, *light blue*) were also plotted but are not visible because no significant edge differences were identified. Analyses used BDBNs from the BN50 (A), BN25 (B), and BN5 (C) pairs. D: Mean number of significant differences in edge frequencies identified between sets of 100 BDBNs corresponding to synthetic BNs differing from the original by varying proportions of edges. A 0% underlying difference represents AD-W-BN comparisons. E: Minimum number of BDBNs required for B-BN or AD-W-BN comparisons to identify significantly more edge differences than SD-W-BN comparisons. *Dark blue values* on bars indicate bar heights. All bars show mean results from 10 repeats; error bars indicate ±1 standard error across the 10 repeats. F: Mean edge differences in BN50 B-BN comparisons (*blue, solid*) and two SD-W-BN comparisons corresponding to each underlying BN (*green, dashed; orange, dotted*) across BDBN set sizes. No significant differences in edge frequencies were identified for any of the SD-W-BN comparisons, meaning both lines lie on the x-axis. *Light blue shading* shows ±1 standard error. G: As in F, but for AD-W-BN comparisons, with each SD-W-BN comparison corresponding to one of the independently simulated dataset.

The ability to detect differences even between very similar underlying networks (BN5) raised concerns that the approach might also identify spurious differences when comparing BDBNs derived from independent datasets representing the same underlying system. Indeed, across-dataset W-BN (AD-W-BN) comparisons also yielded significant differences, indicating that minor dataset variation can produce FPs when comparing B-BN edge frequency differences to SD-W-BN (Figs 5A-C). However, significantly more differences in edge frequencies were identified for B-BN than for AD-W-BN comparisons for all BN50 repeats and 8/10 BN25 and BN5 repeats (S8 Fig). This suggests that using AD-W-BN comparisons as a baseline for the level of spurious differences expected when no true differences exist may be a more robust approach, even for detecting very subtle differences.

Notably, the number of significant edge differences declined as the similarity between underlying networks increased, suggesting this approach may also provide some limited capacity to detect the extent of structural differences (Fig 5D).

The ability to detect smaller differences between underlying networks also depended on the number of BDBNs compared. With fewer bootstraps, significant differences were progressively lost until none were detectable (Figs 5F,5G; S8 Fig). The minimum number of BDBNs required to detect significant differences increased as networks became more similar: only modestly from BN50 to BN5 (23.20±2.99 vs. 28.00±9.12), but much more substantially for AD-W-BN comparisons (44.40±18.37; Fig 5E). This suggests that the minimum number of BDBNs required to detect significant differences may serve as an indicator of whether underlying networks differ, but is unlikely to provide sufficient resolution to assess the extent of that difference.

### Network differences based on DeltaCon similarity scores

Edge frequency comparisons were able to identify that networks differed; however, the edges identified as different did not correspond to true positive edge differences. Given this ended up being a whole-network comparison after all, the analysis shifted from comparing individual edges to assessing global network connectivity using DeltaCon similarity scores. For the BN50 pairs BDBNs showed small but significant reductions in mean DeltaCon similarity scores for B-BN comparisons relative to both SD-W-BN and AD-W-BN comparisons (*p <* 0.0001 across all ten repeats; Fig 6A; S10 FigA).

**Fig 6.**
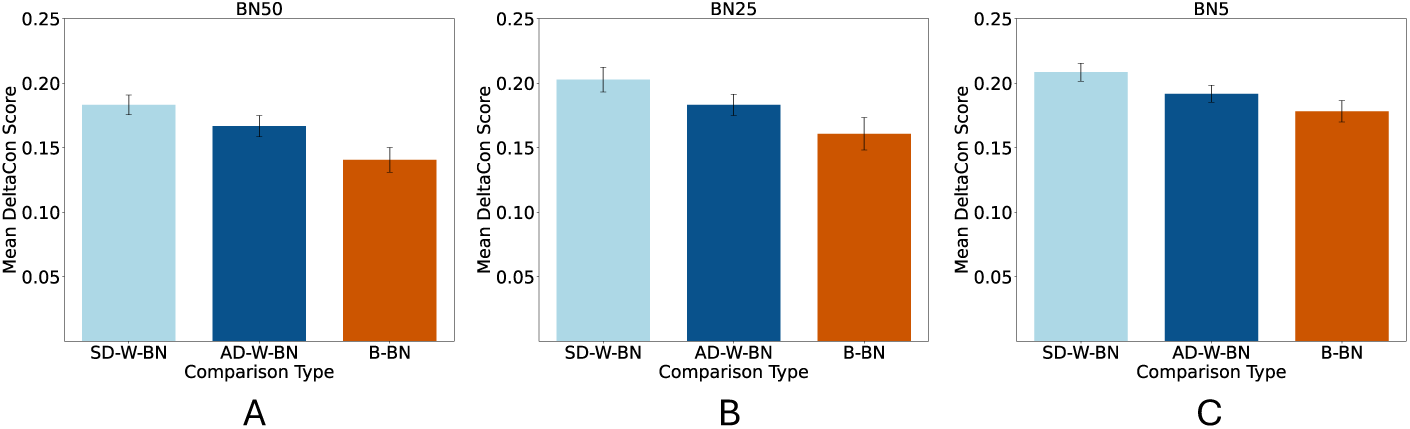
DeltaCon scores between sets of bootstrap-derived Bayesian network (BDBN) Mean number DeltaCon similarity scores between sets of 100 BDBNs from same-dataset within-BN (SD-W-BN, *light blue*), across-dataset within-BN (AD-W-BN, *dark blue*), and between-BN (B-BN, *orange*) comparisons. Analyses used BDBNs from the BN50 (A), BN25 (B), and BN5 (C) pairs. Error bars indicate ±1 standard error across the 10 repeats.

To test the sensitivity of the DeltaCon approach to detecting smaller underlying differences, the same analysis was conduced for the BN25 and BN5 pairs. In both cases, B-BN comparisons showed significantly lower similarity scores than SD-W-BN comparisons in all ten repeats (*p <* 0.0001), while B-BN scores were significantly lower than AD-W-BN scores in 8/10 and 9/10 repeats for the BN25 and BN5 pairs, respectively (Figs 6B-C; S10 FigB-C).

## Discussion

This study developed two bootstrap-based criteria for comparing BNs: edge frequency comparisons and whole-network connectivity. For both approaches, we show that even networks differing by only a few edges (∼5%) can be distinguished as significantly different using relatively small datasets (40 samples). However, reliably identifying the specific edge differences proved much more difficult. Our findings highlight the challenges of distinguishing specific structural differences between networks.

Nevertheless, these criteria provide a framework for more standardised comparisons of BNs that is less sensitive to spurious variation arising from dataset noise or stochasticity in network learning. They have potential applicability across diverse real-world contexts where distinguishing genuine structural differences from sampling artefacts is essential. For substantially different underlying networks (∼ 50%) both the edge frequency and DeltaCon criteria were able to consistently distinguish differences between sets of BDBNs taken from different systems from spurious differences arising from variation in datasets taken from the same underlying network. However, this distinction became less consistent for smaller differences (∼ 25% and ∼ 5%). The similar performance of the edge frequency and DeltaCon criteria may reflect that both, in different ways, capture aspects of indirect as well as direct connectivity. The whole-network DeltaCon approach captures small changes in path lengths, providing a more nuanced basis for assessing differences between BNs. Whilst individual edge comparisons between networks would treat such alterations as entirely distinct, comparing edge frequencies across sets of BDBNs may implicitly reflect indirect relationships. This is because variation across BDBNs can capture alternative edge configurations connecting correlated nodes, thereby integrating information about indirect dependencies into the edge frequency distribution.

While both criteria effectively distinguished when underlying networks differed, the interpretability of their outputs warrants caution. For edge-based comparisons, the number of significant edge differences between sets of BDBNs does not correspond directly to the number of structural differences between the underlying networks. Moreover, the presence of non-zero edge differences in AD-W-BN comparisons, although generally significantly lower than in B-BN comparisons, highlights that dataset variability alone can produce apparent differences. In contrast, DeltaCon similarity scores provide a more abstract but less easily misinterpreted measure of overall structural similarity.

Notably, neither approach could reliably indicate the extent of difference between underlying networks, which remains an important direction for future work. Moreover, both approaches were unable to reliably recover true edge differences between underlying networks. DeltaCon detects overall structural differences but not which edges differ, while edge frequency comparisons identified edges that did not consistently correspond to the true underlying differences, yielding only a handful of TP differences and a large number of FPs. Contrary to what may have been expected, this remained the case even with substantially larger datasets with thousands of data points. Thus, while the number of significantly different edges provides a useful signal to indicate whether underlying network processes producing the data are different, the specific edges do not reliably reflect network structure differences. These findings are consistent with previous observations that edges identified from BDBNs are less consistently recovered than higher-level network features [9]. Therefore, to investigate specific aspects of network differences, alternative approaches could instead compare node centrality distances, which provide higher-level summaries of node roles within the networks [17]. For instance, in BNs representing GRNs, comparing degree centrality distance could highlight differences in hub genes, betweenness centrality distance could indicate changes in important intermediary genes, and closeness centrality distance could reveal broader shifts in how influential particular gene clusters are.

It is important to note that, in real-world applications, AD-W-BN comparisons would require two independent datasets for each condition. Where this is not feasible, each dataset would need to be halved and bootstrapped separately to mimic independence, which in turn necessitates larger overall sample sizes. Since low data availability is often unavoidable, particularly in non-model organisms, the small datasets used here were chosen to represent a particularly stringent test case for the criteria. The relatively large number of nodes relative to sample size further compounded this challenge, meaning that if the criteria performed well under these stringent conditions, they would likely generalise effectively to analyses with larger datasets or simpler networks. Although increasing dataset size did not improve the ability of edge frequency comparisons to identify true edge differences, both criteria’s consistent broader ability to distinguish whether underlying networks genuinely differ demonstrates that both criteria remain robust even with limited data. Nevertheless, dataset size may have exacerbated the inability of bootstrapped datasets to fully recapitulate the variability of independent datasets. Future work should therefore repeat these analyses across a range of network sizes, dataset sizes, and proportions of structural difference between underlying networks to more fully assess the robustness and generalisability of the criteria. Throughout this study, non-parametric bootstrapping was applied, resampling directly from the original dataset. Alternatively, parametric bootstrapping, simulating data from a BN learned from the original dataset, could have increased variability between bootstrapped replicates [28]. While previous research reported limited benefit from parametric bootstrapping at sample sizes of 100 [9], it may have been more advantageous in this case given the smaller datasets. However, the effectiveness of this approach would depend heavily on the quality of the initial BN and the edge probability threshold applied, with lower thresholds allowing more edges and potentially greater variation in simulated samples. Further research could also compare the effectiveness of parametric and non-parametric bootstrapping for smaller datasets.

The edges included in BNs throughout this project were highly dependent on the edge probability threshold applied. While the higher 0.75 edge probability threshold improved network recovery precision relative to 0.5, it inevitably excluded weaker yet potentially still informative edges. Incorporating edge probabilities as weights, rather than applying hard thresholds, could retain valuable information about connection strength. This would complicate edge frequency comparisons but could be readily integrated into computing DeltaCon scores by encoding weights into the network affinity matrices, in place of binary values [19]. Weighted edges could improve sensitivity to subtle network differences and enhance identification of hub clusters by considering the cumulative weights of node connections. Applying a soft-threshold to these weights, as implemented in WGCNA, could promote scale-free topology, enhancing the biological plausibility of the networks by emphasising stronger connections while retaining weaker ones [29, 30].

Unlike in real-world applications, this study’s use of datasets simulated from synthetic BNs provided a known ground truth against which the criteria could be validated. However, random synthetic networks may not fully capture the properties of biological systems. To strengthen the validation, future work should apply the criteria to well-characterised real-world networks. For example, applying them to model organisms with established changes in gene regulatory networks across conditions could provide more robust validation in biologically relevant settings [31, 32].

## Conclusion

Together, these findings demonstrate the potential of bootstrap-derived criteria for comparing BNs using either edge-based or whole-network comparisons. Both approaches can reliably identify when underlying networks of interactions differ, even when these differences involve as little as 5% of edges. This establishes a foundation for a more standardised framework for comparing BNs, while also highlighting the challenges of pinpointing individual edge differences and the risk of sample variation producing spurious differences. Future work should prioritise validating these criteria across a wider range of scenarios and incorporating weighted network approaches to improve sensitivity to smaller structural differences and robustness against sampling noise.

## Acknowledgments

We thank Dr Carmel McDougall for her ideas and advice throughout this research.

## Supporting information

### S1 Appendix Types of heuristic searches

The two main types of heuristic searches are greedy search and simulated annealing (SA), which differ in how they determine whether to accept a new network. Greedy searches only accept higher-scoring BNs and therefore quickly identify local maxima. This is useful for simple search spaces where one structure fits the data substantially better than others. However, in more complex search spaces, it can become trapped in local maxima, preventing discovery of the globally optimal network [3, 33].

In contrast, SA always accepts higher-scoring networks but also has a probability of accepting lower-scoring ones. This probability depends on the “temperature” parameter, which decreases over iterations according to a predefined “cooling schedule”. Early in the search, the higher temperature allows more exploration by accepting worse solutions, while later, as the temperature decreases, the algorithm behaves more like a greedy search. Although slower, SA is better suited for complex search spaces as it reduces the risk of getting trapped at local maxima [33, 34].

Several aspects of the search process can be fine-tuned. For instance, the network modification stage typically involves making a single “local move” by adding, removing, or reversing an edge. The search can either apply one random local move per iteration or evaluate all possible local moves and select the highest scoring one [35]. The latter results in faster searches but is more deterministic. Furthermore, to reduce the risk of getting trapped at local maxima, searches can be configured to restart multiple times within the same run. Stopping criteria, such as maximum number of restarts or networks evaluated, can also be specified [8, 35]. The optimal search settings should be selected on a case-by-case basis to suit the search space characteristics [36].

### S2 Appendix. Preliminary explorations of network search space complexity

To inform the design of a reproducible BN search procedure, an initial qualitative analysis of search-space complexity was conducted by comparing the outdegree consistency of top networks and edge consistencies of consensus networks (constructed from the top 10 and 100 networks) across sets of five repeated greedy and SA searches. Here, outdegree refers to the number of children each node has. This was repeated with stopping criteria of 1 billion and 10 billion network evaluations per search to assess how search length affected consistency. These searches were performed on simulated data.

Greedy heuristic searches evaluating 1 billion networks showed low consistency in the top 10 networks, both within a single search and across repeated searches. Using SA improved within-search consistency, but between-search consistency remained low (S1 FigA-S1 FigB). Increasing the number of networks visited per search to 10 billion only slightly improved consistency (S1 FigC) but substantially increased runtime, from ∼ 2 hours to ≥20 hours, making further increases infeasible. There was also low edge consistency in Banjo-computed consensus networks across searches when computed from both the top 10 and top 100 scoring networks in each search (S1 FigD). Together, these findings suggested that the network search space was complex with many local maxima.

**S1 Fig.**
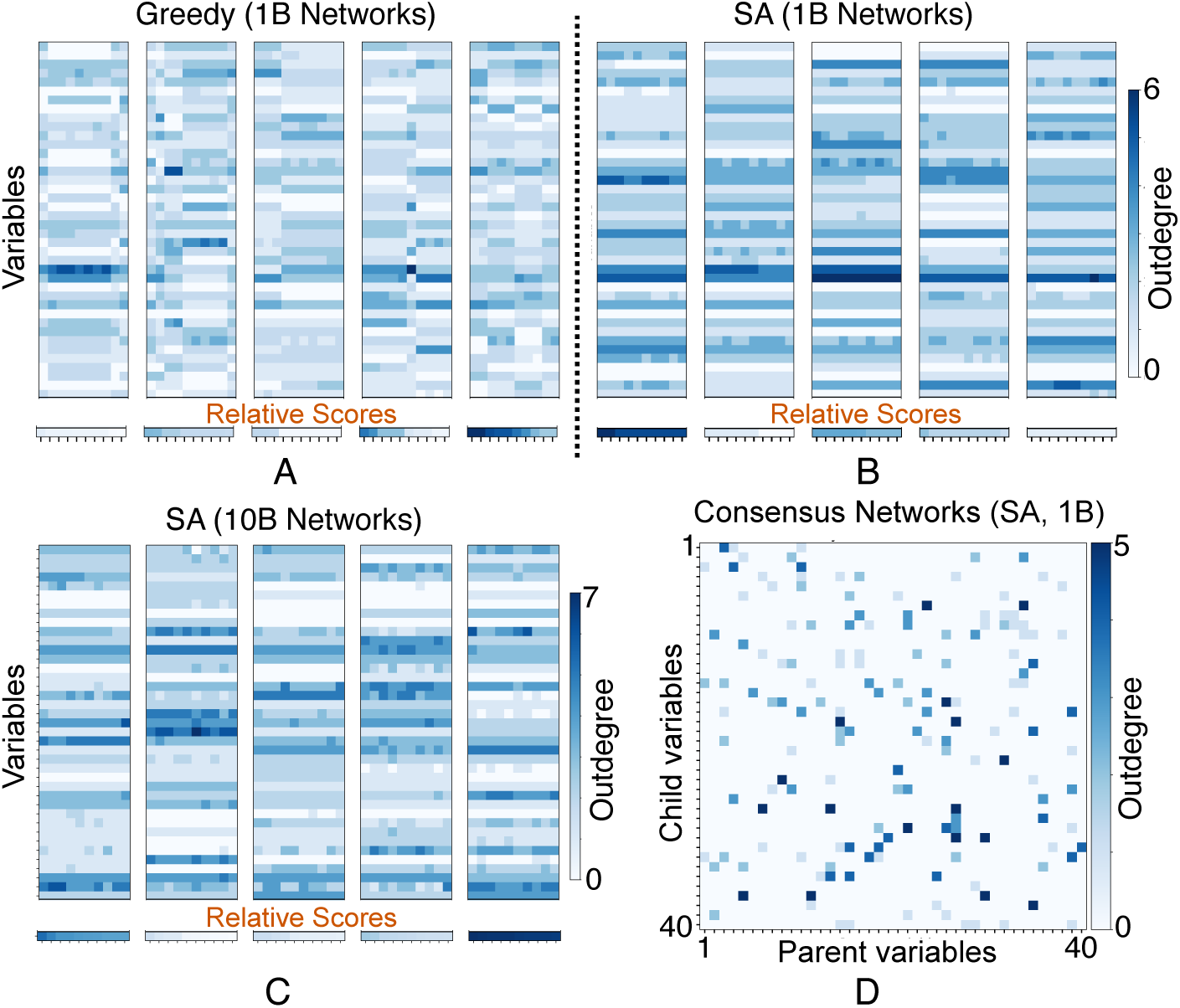
Initial search space explorations. A-C: Outdegree consistency across searches. The outdegree (*blue intensity*) of each variable (*rows*) is shown for the top 10 networks (*columns*) from 5 independent search repeats (*blocks*) using greedy search (A) and simulated annealing (SA; B, C), each evaluating either 1 billion networks (A, B) or 10 billion networks (C). SA shows better within-search consistency (*horizontal stripes*), but both search types have poor between-search consistency (different patterns between blocks). Increasing the number of network evaluations per search provides negligible consistency improvements. The *bottom row* in each block shows relative network scores, with *darker shades* indicating better scores. D: Edge consistency (*blue intensity*) in Banjo consensus networks from 5 independent SA searches evaluating 1 billion networks each. Each consensus network was computed from the searches’ top 10 networks. More *pale blue* indicates lower consistency.

### S3 Appendix. Heuristic search optimisation

To address the complex search space, a network averaging approach was adopted, combining the top scoring networks from 100 greedy searches (See Materials and Methods, S2 Fig). This approach achieved high consistency in edge probabilities between repeated searches (S3 FigA).

Given the large number of searches required throughout this study, a faster stopping criterion than used for initial search space explorations (S2 Appendix) was necessary to ensure feasible runtimes. Following the approach of [7], this criterion was based on the number of restarts, defined as instances where the search encounters a local maximum and restarts from a new random network. Restart limits of 1, 10, 100, and 1000 were trialled, and edge probability variability and the ability to recover edges from synthetic BNs was compared across searches using these different stopping criteria.

The variation in the edge probabilities between searches was assessed to evaluate their impact on reproducibility. For each trial, variability was summarised by calculating the standard deviation (std) of each edge’s probability across 10 repeated search procedures, then averaging these values across all edges.

The variability decreased slightly when increasing the number of restarts per search from 1 to 100, but no further reduction was observed at 1000 restarts (S3 FigB). The largest drop in variability occurred between 1 and 10 restarts, with only marginal improvements beyond this point.

The ability of these search procedures to recover edges in true underlying networks was assessed using a dataset simulated from a synthetic network. After applying a minimum edge probability threshold of either 0.5 or 0.75, edges in each inferred network were compared to the known ground-truth edges from the synthetic network, and the number of true positives (TP), false positives (FP), and false negatives (FN) was counted (S3 FigC). From these, the precision, recall, and F1 score were calculated:

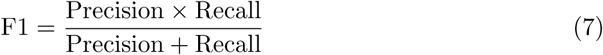

The F1 score summarises the overall ability of a search procedure to recover the underlying network structure, with values ranging from 0 to 1 and F1=1 indicating perfect reconstruction. The mean and std of precision, recall, and F1 were then calculated across the 10 repeated search procedures.

Recall and F1 scores increased with more restarts, while precision improved between 1 and 10 restarts but showed little change beyond that (S3 FigD). Given the diminishing returns and substantially increased runtimes at higher restart limits, a stopping criterion of 10 restarts was selected. Lowering the probability threshold from 0.75 to 0.5 increased recall but reduced precision, due to fewer FNs and more FPs, respectively. For both thresholds, precision remained substantially higher than recall. Since the primary aim of this study was to identify true differences between networks, precision was prioritised over recall. Therefore, a minimum edge probability threshold of 0.75 was chosen.

**S2 Fig.**
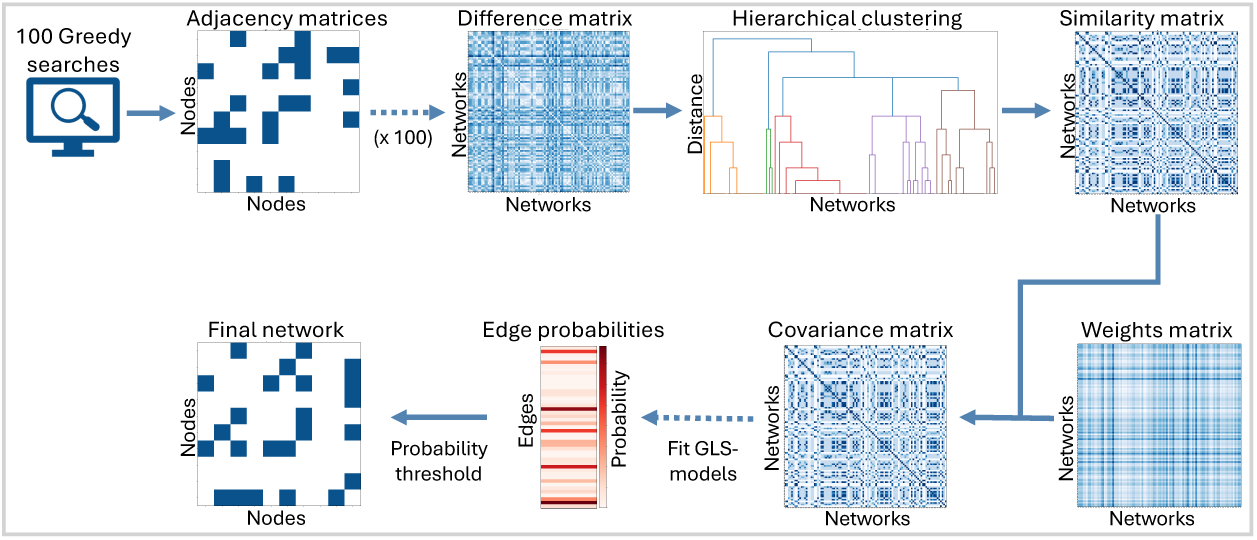
Network averaging search procedure. Workflow for network averaging (10 node example). A difference matrix is computed from the adjacency matrix from 100 greedy searches. Clustering is applied to this to generate a similarity matrix which is combined with a weights matrix, representing network scores, to form a covariance matrix. The covariance matrix is used to fit a GLS-model for each edge to compute its probability of being in the network. A probability threshold is applied to determine which edges are included in the final network.

**S3 Fig.**
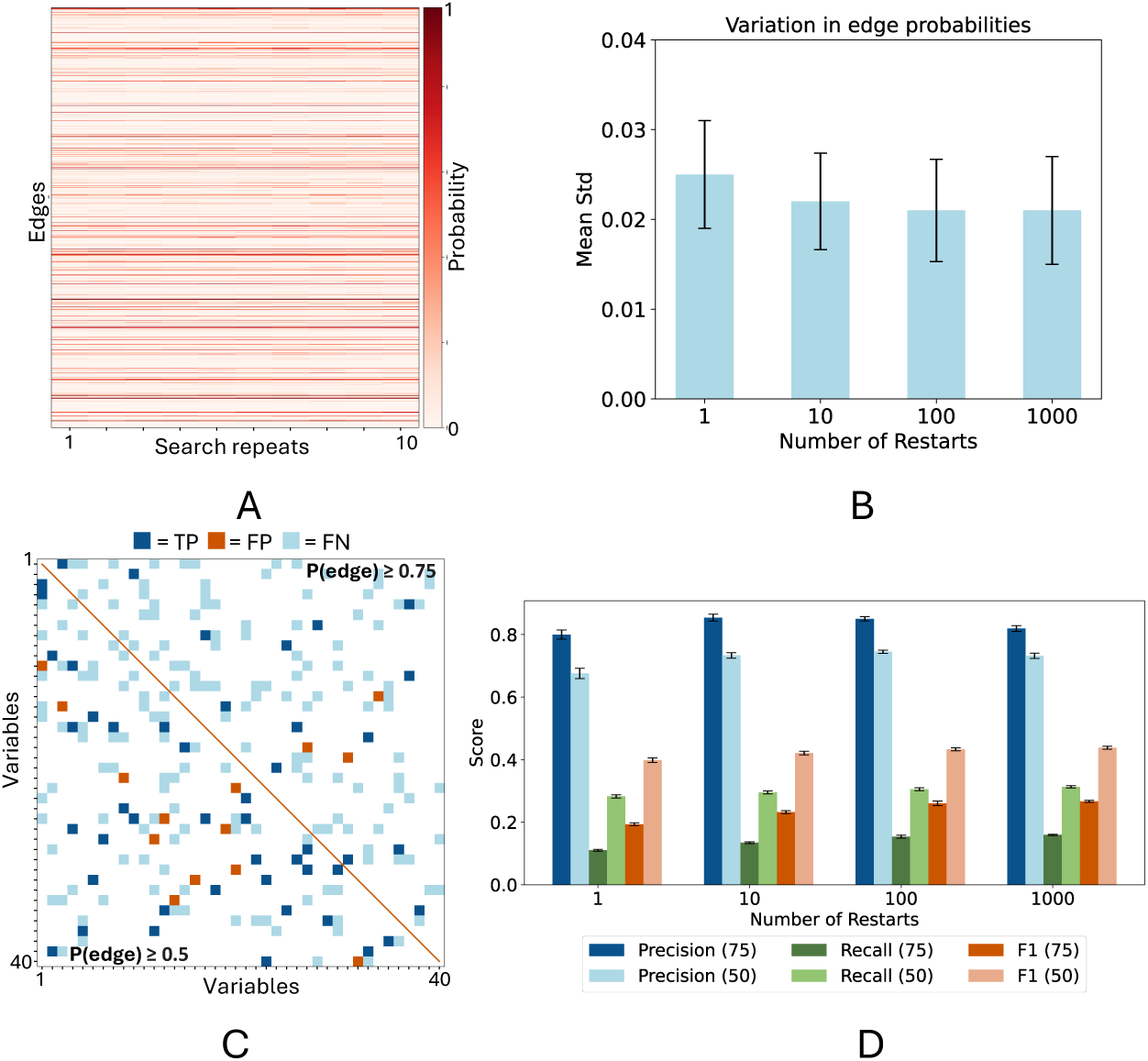
Search reproducibility. A: Example of probabilities (*red intensity*) of possible edges (*rows*) across 10 repeated searches (*columns*). Horisontal stripes of similar shades represent high consistency B: Variation in edge probabilities across 10 repeats from search procedures on simulated data in which each individual search run used varying numbers of restarts (1-1000). Variation was calculated as the mean of the standard deviations (std) in edge frequencies across each individual edge. C: Example recovery of a synthetic Bayesian network (BN) using network averaging, applying edge probability thresholds of 0.75 (*top right half*) and 0.5 (*bottom right half*). True positives (TP, *dark blue*), false positives (FP, *orange*), and false negatives (FN, *light blue*) represent edges correctly recovered, incorrectly added, and missed, respectively, when comparing the learned BN to the synthetic BN. D: Mean precision (*blue*), recall (*green*), and F1 scores (*orange*) for recovering edges in inferred BNs compared to the ground-truth synthetic network across 10 repeated search procedures. Search procedures used varying numbers of random restarts per search (1–1000) and applied a minimum edge probability threshold of either 0.75 (*darker bars*) or 0.5 (*lighter bars*). Error bars in C and D indicate ±1 standard error across 10 repeats.

**S4 Fig.**
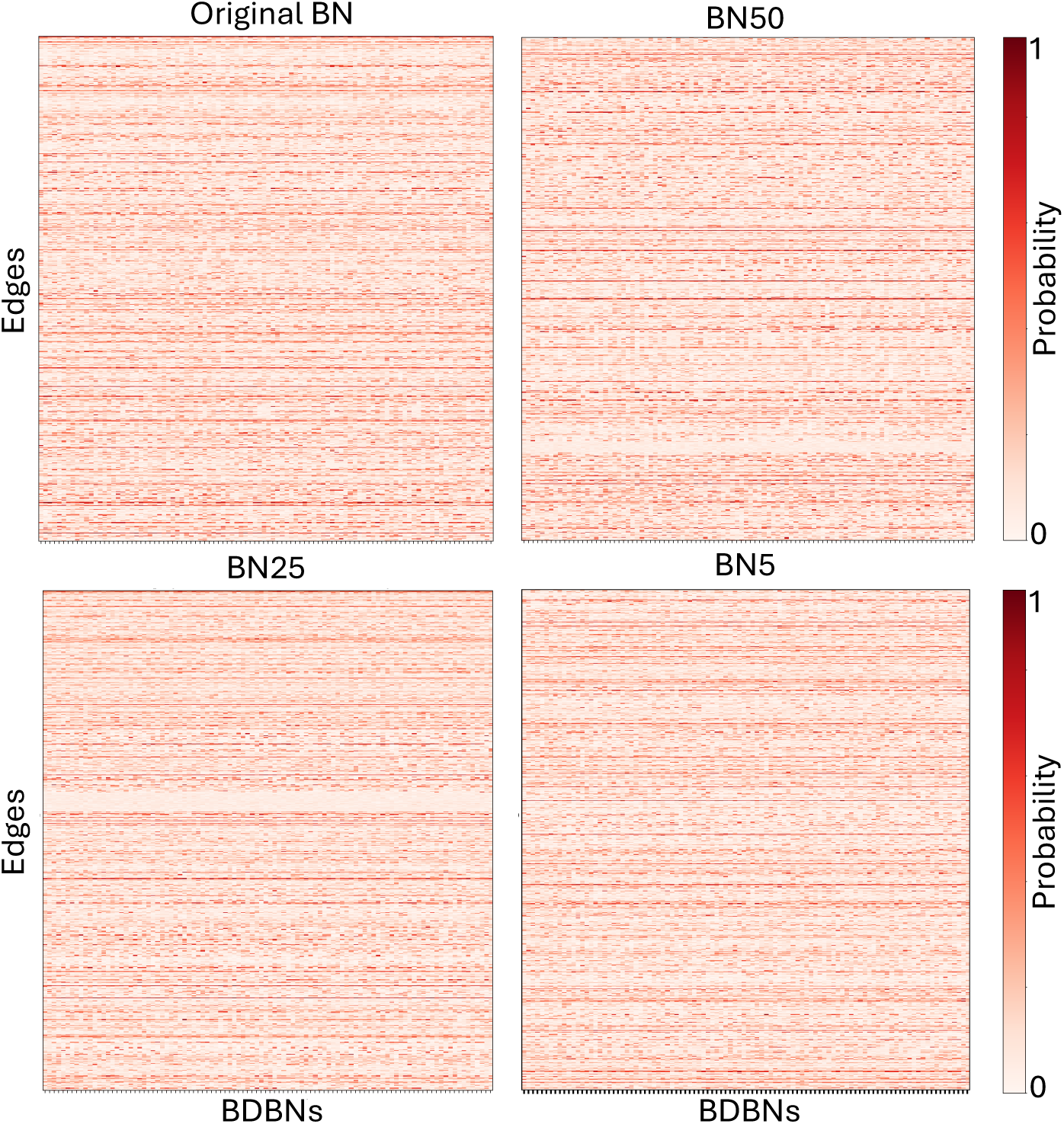
Edge probabilities across bootstrap-derived networks. The probabilities (*red intensity*) of possible edges (*rows*) from example sets of 100 bootstrap-derived Bayesian networks (BDBNs, *columns*) corresponding to either the original randomly generated synthetic BN (*top left*) or the BNs produced by altering either ∼ 50% (BN50; *top right*), ∼ 25% (BN25; *bottom left*), or ∼ 5% (BN5; *bottom right*) of its edges. Horizontal stripes of similar shades indicate consistency across BDBNs from the same set, whilst differences between blocks indicate variation in edges between sets.

**S5 Fig.**
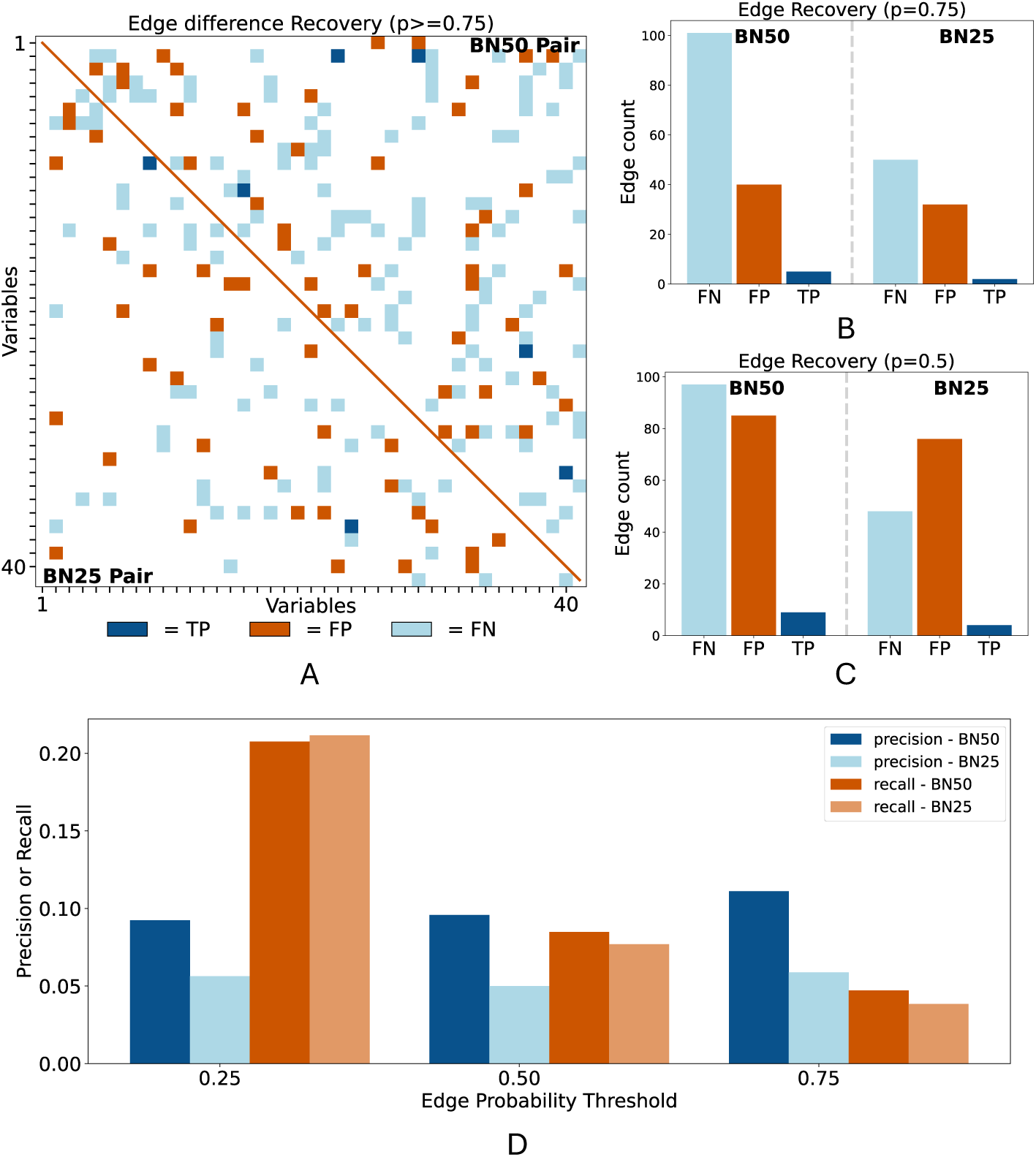
Recovery of differences using datasets of 400 samples. Example of recovering true differences between underlying networks from bootstrap-derived Bayesian networks (BDBNs) learned from 400-sample datasets. A: Recovered differences between the BN50 (*top right*) and BN25 (*bottom left*) network pairs after applying a minimum edge probability threshold of 0.75.True positives (TP, *dark blue*), false positives (FP, *orange*), and false negatives (FN, *light blue*) represent edges correctly identified as different, incorrectly identified as different, and incorrectly identified as not different, respectively. B,C: TP (*dark blue*), FP (*orange*), and FN (*light blue*) counts for BN50 (*left*) and BN25 (*right*) after applying minimum edge probability thresholds of 0.75 (B) or 0.5 (C). D: Precision (*blue*) and recall (*orange*) of edge difference recovery across minimum edge probability thresholds. Results for BN50 and BN25 are shown in *darker* and *lighter* shades, respectively.

**S6 Fig.**
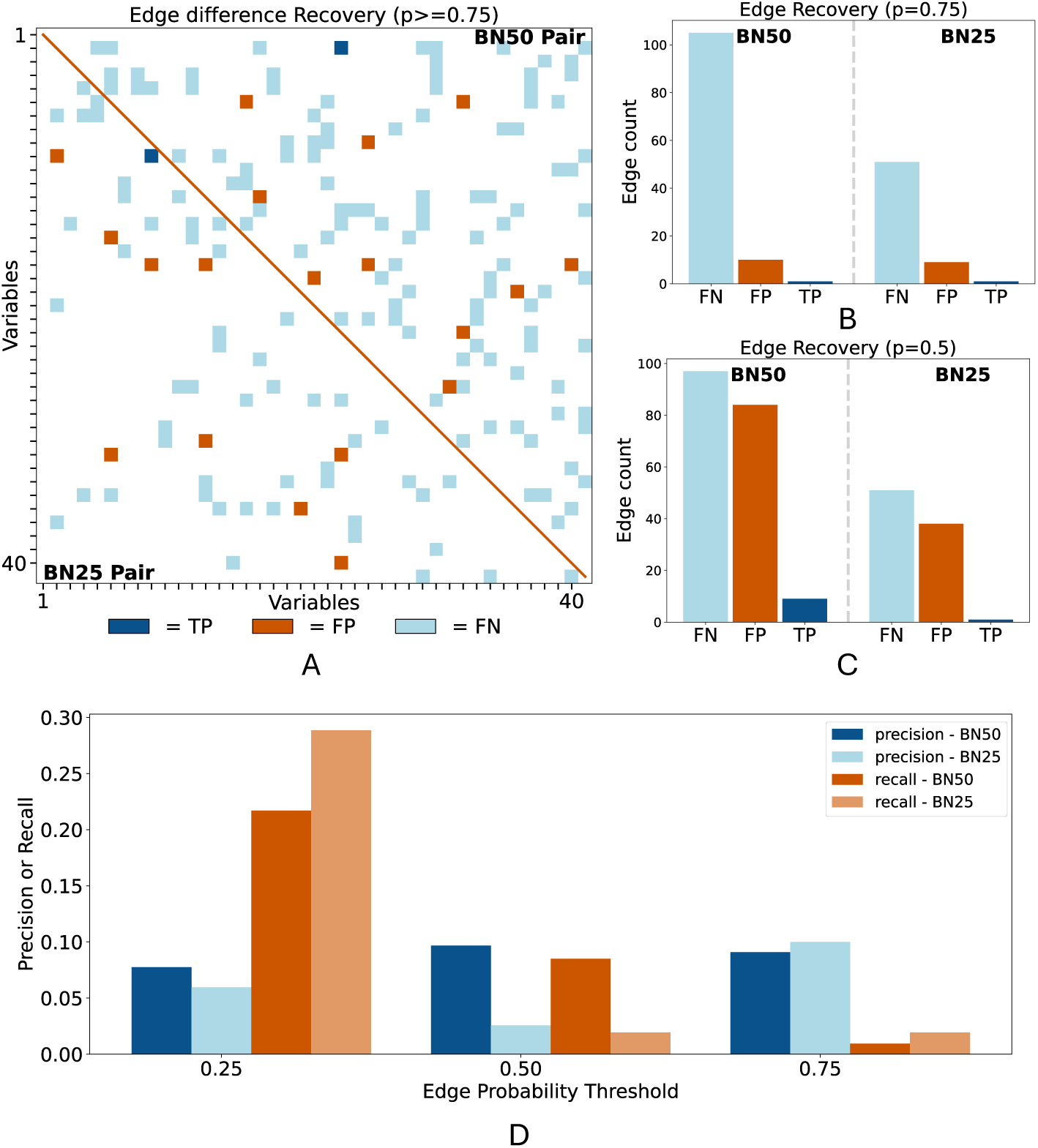
Recovery of differences using datasets of 4000 samples. Example of recovering true differences between underlying networks from bootstrap-derived Bayesian networks (BDBNs) learned from 4000-sample datasets. A: Recovered differences between the BN50 (*top right*) and BN25 (*bottom left*) network pairs after applying a minimum edge probability threshold of 0.75.True positives (TP, *dark blue*), false positives (FP, *orange*), and false negatives (FN, *light blue*) represent edges correctly identified as different, incorrectly identified as different, and incorrectly identified as not different, respectively. B,C: TP (*dark blue*), FP (*orange*), and FN (*light blue*) counts for BN50 (*left*) and BN25 (*right*) after applying minimum edge probability thresholds of 0.75 (B) or 0.5 (C). D: Precision (*blue*) and recall (*orange*) of edge difference recovery across minimum edge probability thresholds. Results for BN50 and BN25 are shown in *darker* and *lighter* shades, respectively.

**S7 Fig.**
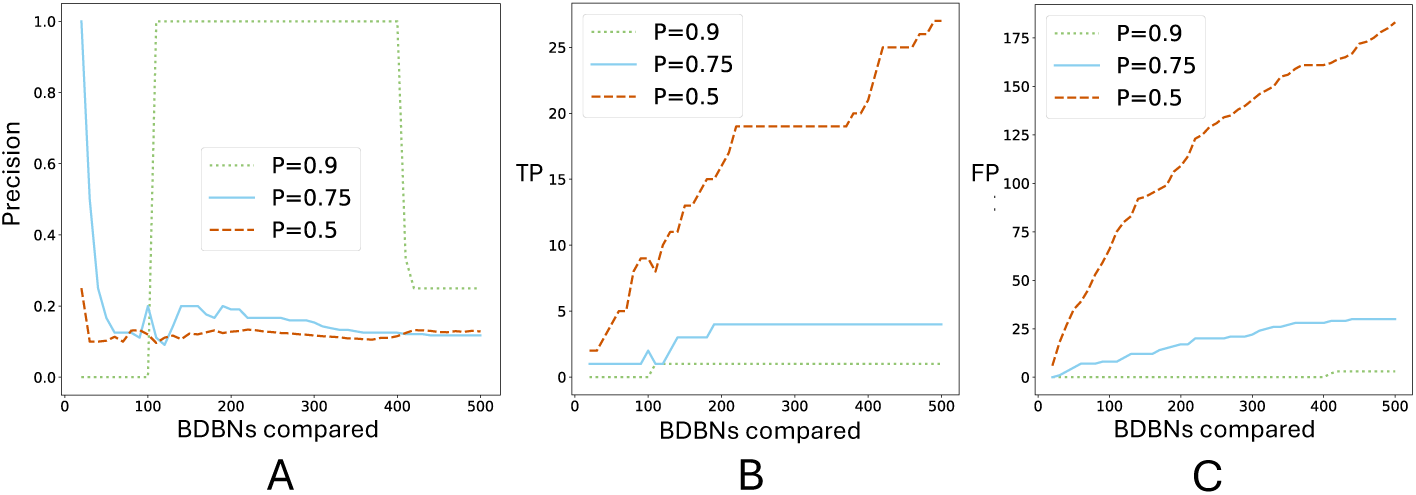
Edge difference recovery across varying probability thresholds and bootstrap-derived networks being compared. Precision in edge difference recovery (A) and number of TP (B) and FP (C) edge differences identified for an example BN50 pair after comparing different numbers of bootstrap-derived Bayesian networks (BDBNs) and applying minimum edge probability thresholds of 0.5 (*orange, dashed*), 0.75 (*light blue, solid*), and 0.9 (*green, dotted*).

**S8 Fig.**
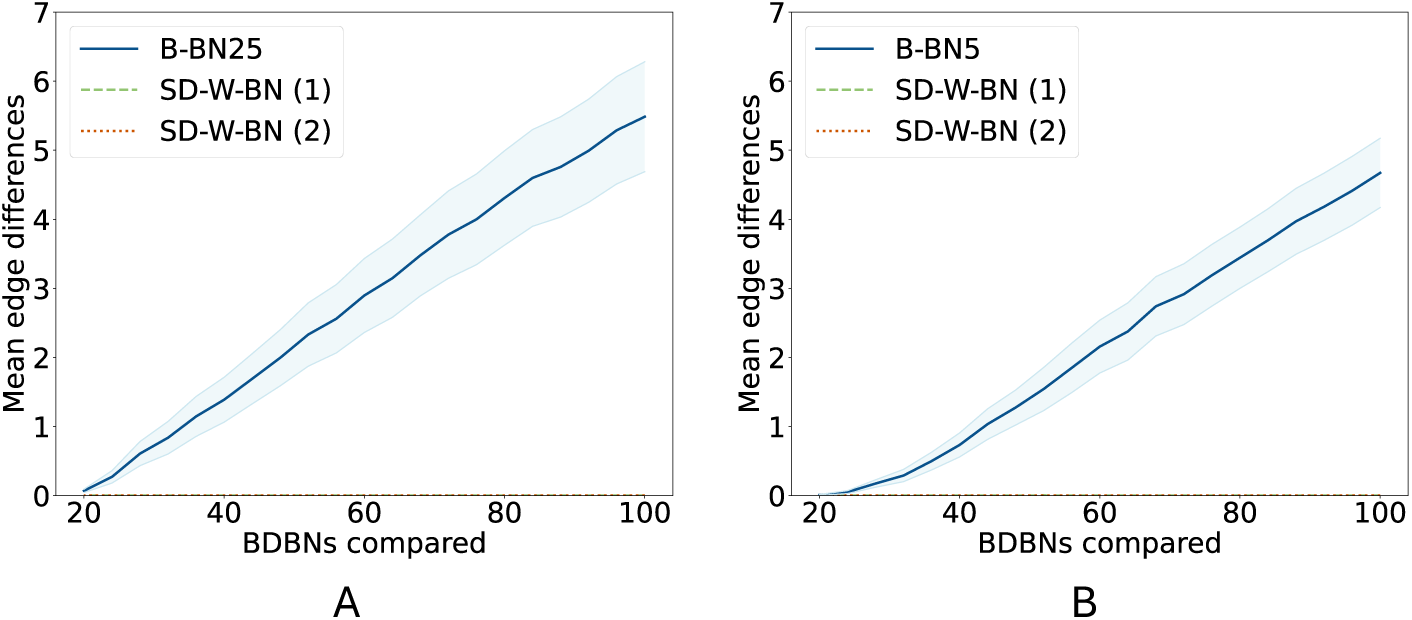
Within- and between-BN edge difference comparisons for smaller underlying differences. Mean edge differences identified in the between-BN (B-BN) comparisons (*blue, solid*) and the two same dataset within-BN (SD-W-BN) comparisons (*green, dashed; orange, dotted*) across varying numbers of bootstrap-derived Bayesian networks (BDBNs). No significant differences in edge frequencies were identified for any of the SD-W-BN comparisons, meaning both lines lie on the x-axis. *Light blue shading* shows ±1 standard error for B-BN comparisons. A: Comparisons of BDNBs corresponding to underlying networks differing in ∼ 25% of edges (BN25). B: Comparisons of BDNBs corresponding to underlying networks differing in ∼ 5% of edges (BN5).

**S8 Fig.**
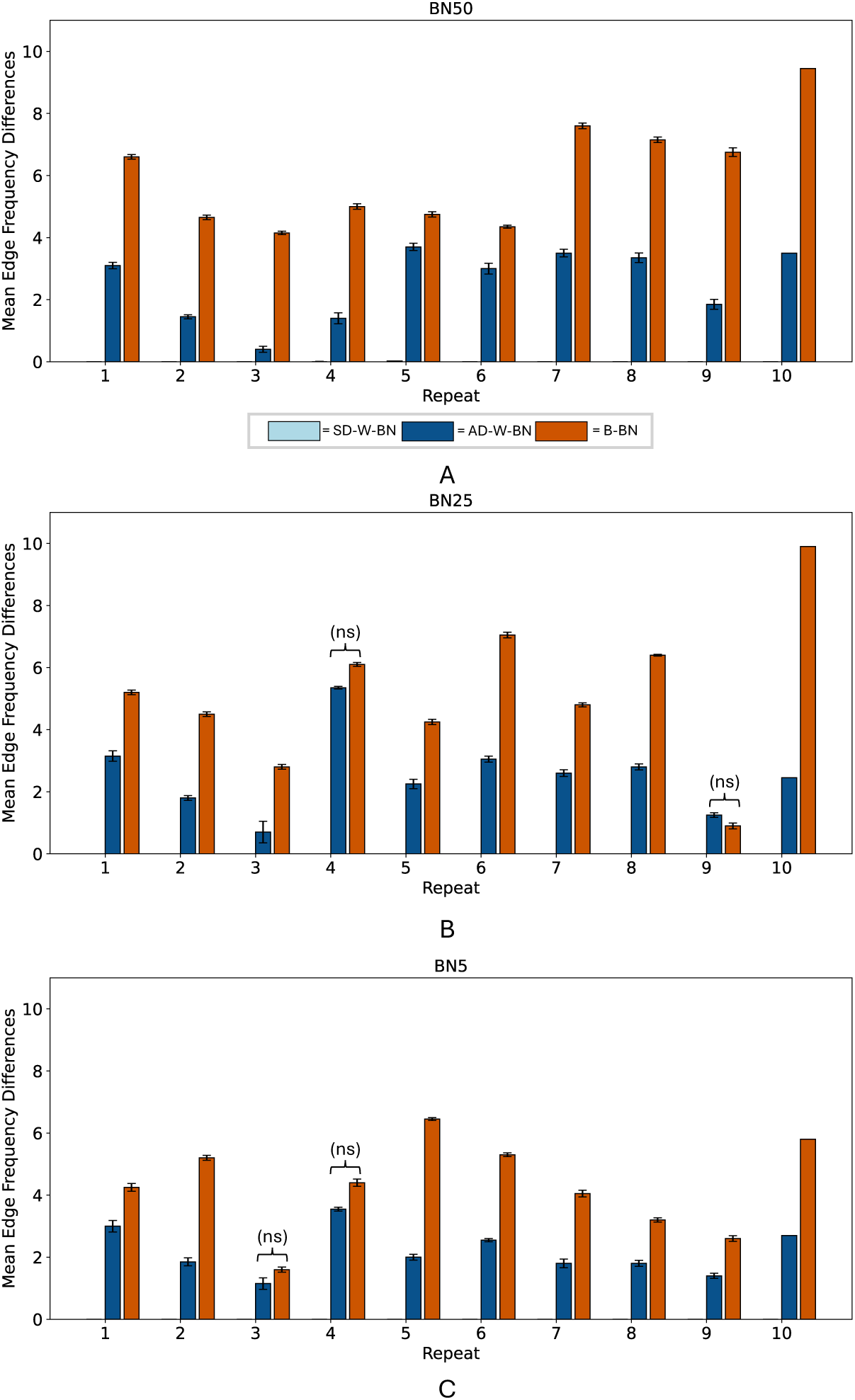
Within- and between-BN Edge frequency differences across repeats. Bars show the mean number of significant edge frequency differences between sets of bootstrap-derived Bayesian networks (BDBNs) across 10 repeats, with error bars indicating ±1 standard error based on 100 random splits of each BDBN set into two halves. Comparisons were made between across-dataset within-BN BDBNs (AD-W-BN, *dark blue*) and between-BN BDBNs (B-BN, *orange*). Same-dataset within-BN (SD-W-BN, *light blue*) comparisons were also plotted but are not visible because no significant edge differences were detected in any repeat. Analyses used BDBNs corresponding to the BN50 (A), BN25 (B), and BN5 (C) pairs. Where indicated, differences between adjacent bars are not significant (ns); all other comparisons are significant at *p <* 0.05.

**S10 Fig.**
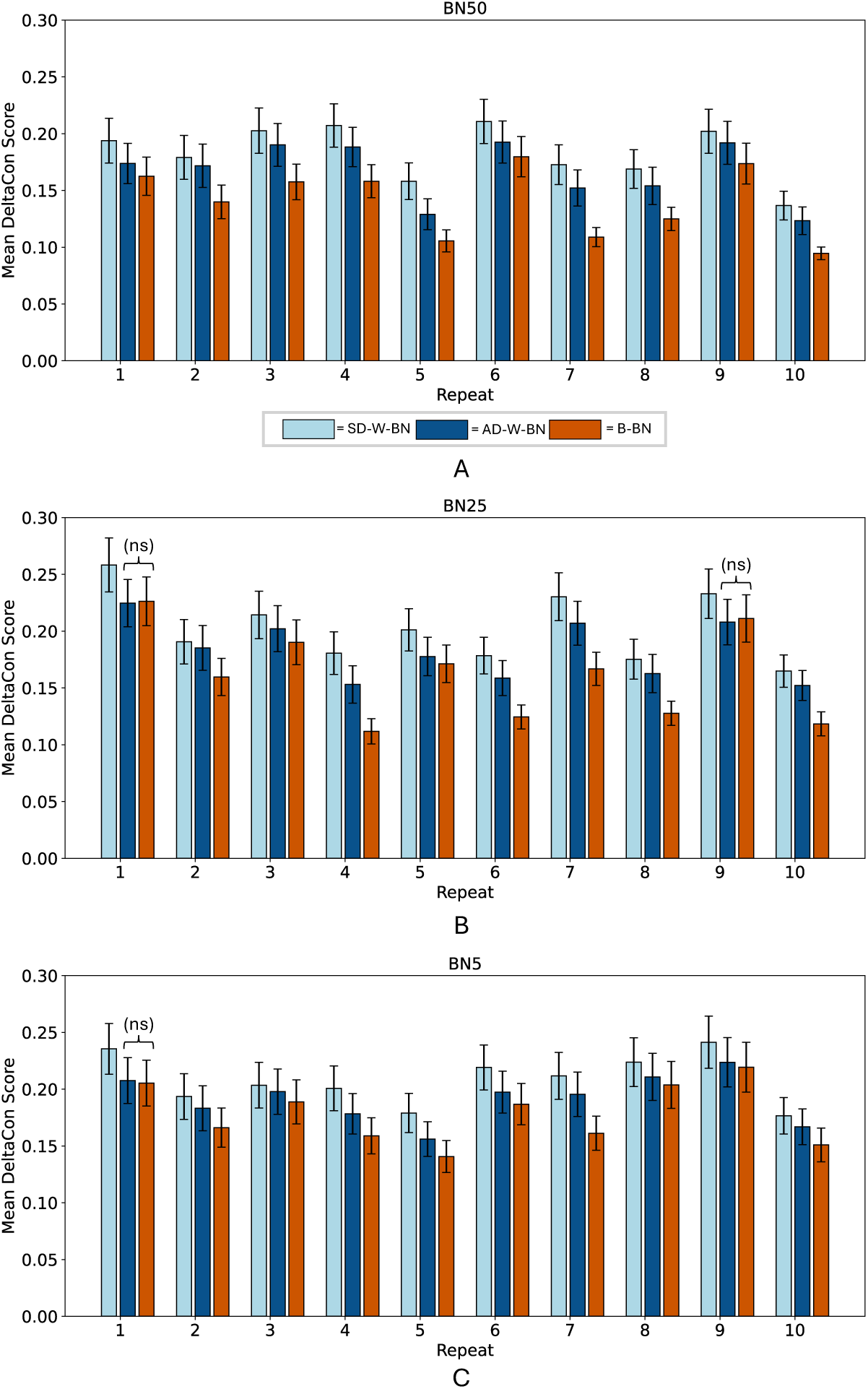
Within- and between-BN DeltaCon comparisons across repeats. DeltaCon similarity scores from same-dataset within-BN (SD-W-BN, *light blue*), across-dataset within-BN (AD-W-BN, *dark blue*), and between-BN (B-BN, *orange*) comparisons. Bars show mean scores for each of 10 repeats; error bars indicate ±1 standard error across 100 bootstrap-derived Bayesian networks (BDBNs). Analyses used BDBNs corresponding to the BN50 (A), BN25 (B), and BN5 (C) pairs. Where indicated, differences between adjacent bars are not significant (ns); all other comparisons are significant at *p <* 0.05.

